# The use of CRISPR for variant specificity in the genetic diagnosis of primary immunodeficiency disease (PID)

**DOI:** 10.1101/804732

**Authors:** Tiffeney Mann, Amy Smith, Sarah Spencer, Alasdair Russell, James Thaventhiran

## Abstract

The functional validation of genetic variants of uncertain significance (VUS) found in PID patients by next-generation sequencing has traditionally been carried out in model systems that are susceptible to artefact. We use CRISPR correction of primary human T lymphocytes to demonstrate that a specific variant in an IL-6R deficient patient is causative for their condition. This methodology can be adapted and used for variant assessment of the heterogeneous genetic defects that affect T lymphocytes in PID.

## INTRODUCTION

Primary immunodeficiency diseases (PIDs) form a diverse group of genetic disorders caused by defects in the development or function of the immune system, which predispose to infectious pathogens but also autoimmune disease and malignancy (1). To date, more than 350 distinct disorders have been described, with a similar number of individual genes implicated (2). Obtaining an accurate diagnosis has important consequences for prognosis, treatment and genetic counselling (3). Traditional PID diagnostic strategies encompass clinical evaluation, and standard laboratory screening tests, as well as testing of specific pathways, for example assays to evaluate neutrophil oxidative burst, or T cell receptor signalling pathways (4). This approach enables clinicians to categorise the patient’s clinical and immunological phenotype and identify one or more candidate genes for Sanger sequencing (1)(5). This phenotype-based approach comes with a significant time and resource cost. It is dependent on obtaining viable cells from patients and training laboratory staff to perform a diverse range of techniques. Transfer of patient samples can impact cell viability and impair physiological responses, compromising the accuracy of diagnostic tests (1). Alternatively, next-generation sequencing (NGS) methods are becoming increasingly accessible in the clinical laboratory setting. These rapid, accurate, and relatively low cost methods allow a high-throughput, genotype-based approach to molecular diagnosis (5). Whole genome sequencing (WGS) has been able to demonstrate the broad range of phenotypes caused by monogenic mutations and has facilitated discovery of multigenic PIDS and risk factor genes/modifier mutations. In addition, high coverage WGS allows identification of somatic variants in subpopulations of immune cells (1).

Next generation sequencing methods do, however, have some drawbacks. Often the approach is to sequence the proband and parents/siblings, to narrow the list of candidate variants. However, elimination of variants present in ‘unaffected’ family members assumes complete disease penetrance, which is not always the case (1). Several software tools have been developed to prioritise variants based on predicted pathogenicity, including CADD, SIFT, PolyPhen2, GERP, GWAWA and MutationTaster. Although powerful, these tools can incorrectly characterise variants and may be inconsistent with each other (6). The huge amount of data generated from NGS, and the vast number of genes sequenced, increases the rate of detection of variants for which clinical significance is uncertain. ‘Variants of uncertain significance’ (VUS) are those which do not meet sufficient criteria for classification as ‘pathogenic’, ‘likely pathogenic’, ‘benign’, or ‘likely benign’. For some VUS, the nature of the variant may be suggestive of pathogenicity (e.g. truncating variant), but it occurs in a gene whose relevance to human disease is not yet known. This is particularly relevant to patients with PIDs, since the pace of discovery in the field is rapid. Secondly, a variant may be classified as a VUS because, despite being identified in a gene relevant to the clinical or immunological phenotype, there is insufficient evidence of pathogenicity (7)(1).

Genetic studies of single patients can be powerful and informative to the field of immunodeficiency. However, such studies are dependent on the meeting of three main criteria in order to confidently attribute phenotype to genotype in a single patient. Firstly, the mutation must be absent in the normal healthy population. Secondly, there must be evidence that the mutation affects expression or function of the gene product. Finally, a causal relationship must be demonstrated by functional studies using a relevant cellular phenotype or animal model (8). Validation of the pathogenicity of mutations may be achieved by transfection experiments: for loss of function mutations, transfection of the wild-type copy of the gene will correct the cellular phenotype, whereas for gain-of-function or dominant-negative mutations, transfection of a mutant copy into wild-type cells will induce the phenotype. However, such experiments are hampered by difficulties with regulation of expression of the transfected gene (8). CRISPR-Cas9 correction of the VUS has the potential to overcome this artefact by correcting the candidate VUS and assessing if function is restored. We have applied this technique to the cells of a patient with recently described transfection-validated IL6R deficiency [ref] and confirmed that correction of the candidate variant to the reference genome restored IL-6R function. The techniques used here could be applied for the assessment of other VUS in genes that affect T lymphocyte function.

## RESULTS AND DISCUSSION

In the recently described IL-6R deficiency, WGS identified uniparental isodisomy of chromosome 1 in a patient with PID (9). Within chromosome 1, a rare frameshifting homozygous mutation ([c.548del] + [c.548del], chromosome 1, exon 4) in the IL-6R gene leading to a premature stop codon and truncated protein was identified. The *IL-6R* is central to IL-6 signalling - IL-6 binding to IL-6R leads to activation of the JAK/STAT signal transduction pathway leading to STAT3-dependent growth and differentiation. The patient cells shown impaired IL-6 induced pSTAT3 and ectopic expression of wild-type IL-6R, restored STAT3 responses to IL-6. However, this experiment does not exclude the possibility that other variants in the highly atypical chromosome 1 of this patient, led to their impaired IL-6R function. Genome editing can precisely determine whether the specific mutation led to the loss of IL-6R function.

A single guide RNA (sgRNA) was designed and assessed computationally for predicted on-target and off-target activity using DESKGEN software package (www.deskgen.com). A 200 bp single-stranded donor template (ssODN) was designed to be homologous with the target DNA 100 bp either side of the mutation. The template contains a 1 bp insertion, designed to correct the patient’s 1 bp deletion of a guanine. Three phosphorothioate (PS) inter-nucleotide linkage modifications, thought to increase HDR efficiency by stabilising the ssODN, were incorporated at the 5’ and 3’ ends with a sulfurizing reagent (10). To circumvent recutting of the corrected patient DNA by excess ribonucleoprotein (RNP), a silent mutation was created in the sgRNA protospacer adjacent motif (PAM) site in the HDR template. T cells (CD3^+^ T cells) were first isolated from the patient’s PBMCs. The sgRNA designed and tested to target *IL-6R* and Spy Cas9 complexed into a ribonucleoprotein (RNP) and the ssODN designed to correct *IL-6R* was electroporated into the patient T cells. Genomic (g)DNA was extracted from half of the cells and Sanger Sequenced (11). The chromatogram from the edited patient sequencing trace in Figure 1 (B) shows superimposed signalling peaks downstream from the cut site, representing a heterogeneous pool of edited sequences. The indel contribution was deconvoluted from the Sanger trace using the Inference of CRISPR Editing tool (12). The inferred sequences present in the population and the relative proportion of each sequence are shown, as are the percentage of alleles in the pool of sequences that incorporated the ssODN through homology directed repair (% HDR). The indel contribution shows the % HDR, here 44%, and the insertion of a G, correcting the patient’s DNA. The silent mutation in the PAM has been incorporated at the same percentage. The other half of the cells were used for functional interrogation using Western Blotting. Whole-cell lysate was prepared, and samples were analysed by Western Blotting using antibodies specific for total STAT3 and phosphorylated (p)STAT3. The data shows restoration of pSTAT3 upon IL-6 stimulation in CRISPR-Cas9 corrected cells compared to the unedited patient control, shown in Figure 1B.

**Figure 1.**
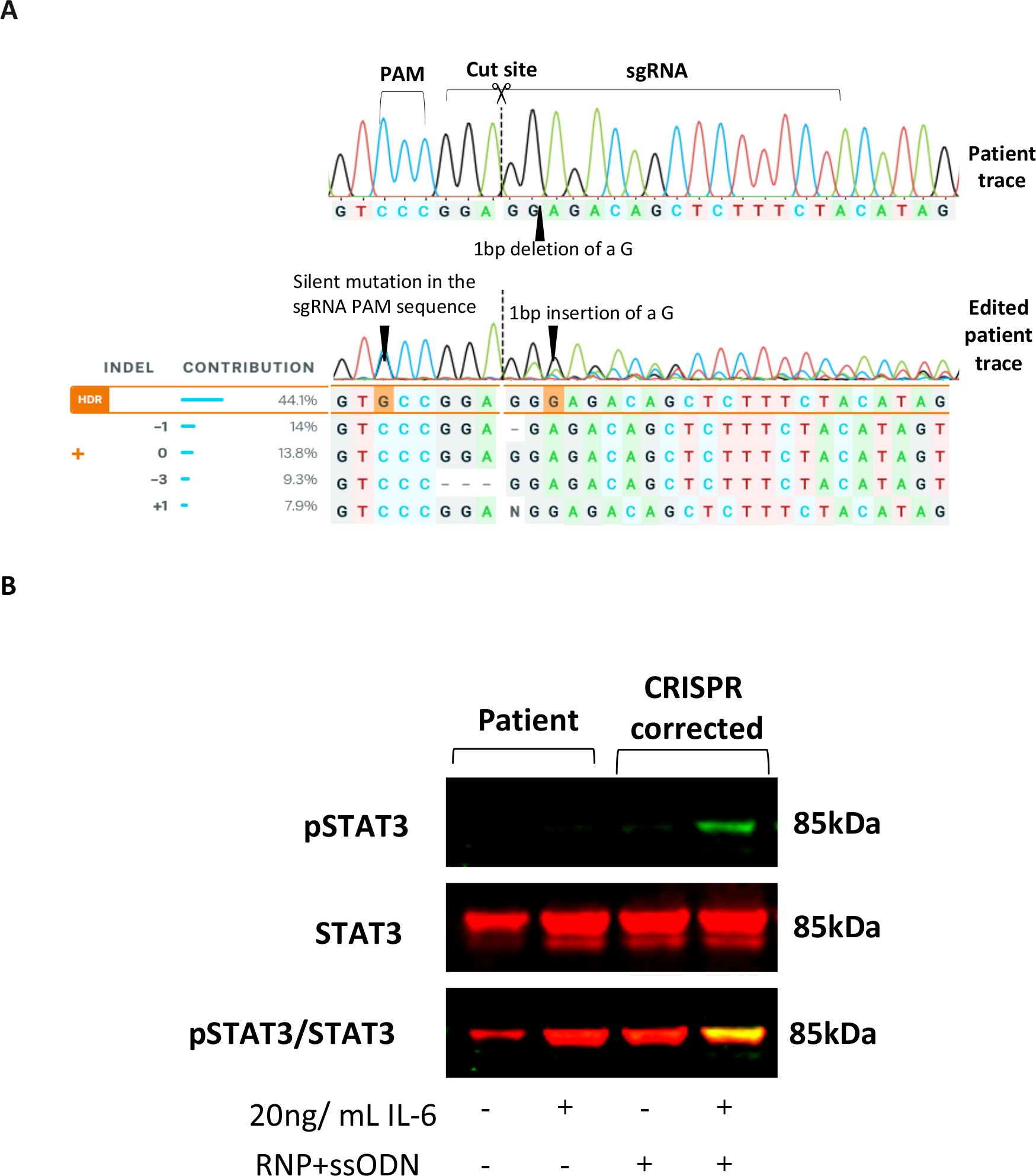
CRISPR correction of patient-derived T cells with an RNP targeting *IL6R* and an ssODN designed to correct the patient’s rare *IL6R* variant and western blot analysis. (a) **Sanger sequencing traces.** The Sanger Sequencing trace spanning the cut site of the unedited patient sample (top) and the edited patient sample (bottom) in this experiment. The black arrow on the edited patient trace shows where the correction (1 bp insertion of a G) has been incorporated. An arrow also points to the position of the silent mutation in the sgRNA PAM sequence. The indel contribution shows the inferred sequences present in the population and the relative proportion of each sequence (example: −1 = 1 bp deletion). The percentage of sequences that have incorporated the single-stranded donor by homology-directed repair (HDR) are boxed in orange. The colour of the peaks represents the base; Black=G, Green=A, Blue=C and Red= T. The black dotted line represents the cut-site. (b) **Western blot analysis of IL-6 induced STAT3 activation in patient and *IL6R* CRISPR corrected patient T cells.** The blot shows STAT3 (red) and pSTAT3 (green) protein expression in patient and CRISPR corrected T cells. Cells were stimulated with IL-6 (20 ng/ mL, 30 minutes). The membrane was probed with both anti-STAT3 and anti-pSTAT3 antibodies, with bands expected to be detected at 85 kDa. Total STAT3 was used as a loading control. RNP= ribonucleoprotein (Cas9 and sgRNA), ssODN= single-stranded oligonucleotide donor. Results from 1 of 2 replicate experiments are presented.

Accompanying the increased application of WGS in clinical diagnostics comes the identification of a growing number of potentially pathogenic genetic variants requiring functional assessment and validation. This CRISPR-Cas9 protocol offers a strategy for rapid assessment and confirmation of causality of VUS in genes involved in signal transduction pathways in immune cells, circumventing viral delivery of CRISPR components. This brings the opportunity to understand the functional impact of genetic change in a relevant model system and it leads to the possibility of therapeutic applications using cell and gene-based therapies. The data here from a model system confirms successful CRISPR-Cas9 correction of a novel genetic anomaly in the *IL-6R* gene and restoration of receptor function, confirming that the patient’s lack of STAT3 response is caused by their rare *IL-6R* variant. Furthermore, the lack of STAT3 signalling caused by the variant helps to explain the clinical phenotype of the patient. Ensuring that the approach is applicable when allele-specific editing is required is key to building this work into a functional genomics platform that is broadly applicable to a range of pathogenic mutations.

Probing genetic function using CRISPR knock-out via the NHEJ pathway is well adopted whereas using CRISPR knock-in via the HDR pathway is less well established and is less commonly utilized due to low efficiency (13). Due to the limited number of patient cells, a biochemical assay was used to detect restoration of STAT3 because this requires fewer cells. Quantifying data using Western Blotting is difficult and comparing editing efficiency from Sanger sequencing data to the functional rescue of *IL6R* is unreliable. Several developments could be made here to enhance the data. A functional single cell-based assay to detect restoration of STAT3 signalling in CRISPR corrected cells would provide more quantifiable results. Single-cell cloning of the edited population of cells could provide paired genotypic and functional information about independently edited clones and downstream experimentation on edited clones such as investigation of transcription factors and expression of genes involved in the inflammatory response. Currently, the data shows the proportion of alleles in the pool of sequences that have been corrected. Introducing genetic markers to purify the cells that have been homogeneously edited would be useful for functional interrogation and potential therapeutic applications.

## MATERIALS AND METHODS

### Cell culture

All cell lines used were authenticated by Short Tandem Repeat (STR) DNA profiling and tested for mycoplasma contamination regularly.

#### Primary human T cells

Human peripheral blood mononuclear cells (PBMC) were isolated by Ficoll gradient centrifugation from a leukocyte reduction system chamber (LRSC) (buffy cone) collected from healthy donors at the NHS Blood and Transplant Centre, Addenbrookes, NHS trust, Cambridge, UK. Written Informed consent was obtained from the patient and their relatives and approved by Cambridge South Research Ethics Committee (13/EE/0325). Whole blood was diluted with PBS supplemented with 0.5 M EDTA and Histopaque 1007 (Sigma-Aldrich) was gently layered over the top. Blood was centrifuged at 800 g for 20 minutes at room temperature with the break set to “off”. The layer of PBMCs was removed using a Pasteur pipette and washed twice using PBS supplemented with 1% FBS, with centrifugation at 400 g for 10 minutes, 4°C. Isolated PBMC’s were resuspended in antibiotic-free RPMI+ Glutamax media (Gibco™), supplemented with 10% heat-inactivated FBS and 0.05 mM of 2-mercaptoethanol (Sigma-Aldrich).

#### PBMC stimulation

A 96 well plate was prepared with 5 μ/mL of purified antimouse-CD3 Antibody (BioLegend, 17A2) diluted with PBS and incubated for 2 hours (37°C, 5% CO_2_). Following incubation, the plate was washed twice with PBS. A cell suspension was prepared at 1×10^6^ cells/mL of RPMI with 5 μ/mL of soluble purified anti-human CD28 antibody (BioLegend, CD28.2) in the presence of recombinant Human IL-2 (carrier-free, BioLegend) and recombinant Human IL-7 (carrier-free, BioLegend) cytokines at 20 ng/mL and 2 ng/mL respectively. 200ul of cell suspension per well was plated into the CD3 bound 96 well plate and incubated for 72 hours at 37°C, 5% CO_2_ to activate CD3^+^ primary human T cells.

### Single guide RNA (sgRNA) and single-stranded oligonucleotide donor (ssODN) design

Target specific sgRNAs were designed using DESKGEN software (www.deskgen.com). DESKGEN software was used with 150 bp’s of patient sequence surrounding the *IL6R* mutation was input into DESKGEN and sgRNA target sites were identified. sgRNAs that overlapped the *IL6R* mutation were prioritised and subsequently assessed for computationally-predicted on-target and off-target activity. On-target activities were calculated using the algorithm of Doench *et al* (14), which predicts how well a given sgRNA will cut the DNA at the desired target site. A score of 100 predicts the highest activity based on the composition of the nucleotide sequence in the sgRNA. To determine off-target activity, the algorithm of Hsu *et al* was used (15). The off-target algorithm predicts how specific a given sgRNA is within a genome of interest. The range of scores varies from 0 to 100, with a higher score predicting that the guide will cut at fewer unintended sites in the genome. The score is based on the similarity of the sgRNA sequence to other sites in the genome and includes sites that have mismatches to the sgRNA. Synthetic Spy Cas9 compatible sgRNAs with 2—O-methyl phosphorothioate linkage modifications at the 3’ and 5’ ends were chemically synthesised (Sigma-Aldrich, Haverhill, UK). IL-6R sgRNA target sequence: 5’-TAGAAAGAGCTGTCTCCTCC-3’.

Lyophilised sgRNAs were resuspended in Milli-Q purified H_2_O to 200 pmol/μL. An ssODN (IDT, 4 nmol ultramer DNA oligos) was designed using the SnapGene® (GSL Biotech LLC) software package and constituted 100 base pair (bp) homology arms around the target modification in *IL-6R.* 2—O-methyl phosphorothioate modifications were designed into the 3 terminal nucleotide linkages at both the 3’ and 5’ ends of the ssODN. The lyophilised ssODN was resuspended in Milli-Q purified H_2_O to 200 pmol/μL.

### Biochemical *in vitro* Cas9 cleavage assay

An *in vitro* Cas9 cleavage assay was used to evaluate the efficiency of sgRNA designs and the effect of the Cas9 nuclease on the editing outcome. Double-stranded 250 bp DNA gBlocks were synthesised against the variant on patient DNA (IDT) as a substrate for cleavage. Two variants of Cas9 nuclease were used in this assay, an evolved highly-specific variant, *eSpy* Cas9 (gift from Dr Andrew Bassett, Wellcome Trust Sanger Institute, Hinxton, UK) and the parental, non-evolved form of *Spy* Cas9 (TrueCut™, *Spy* Cas9, Invitrogen). 200 ng of sgRNA was pre-complexed with 0.5 ng/μL of *spy*Cas9 or *eSpy*Cas9 for 20 minutes at room temperature. 100 ng of gBlock was added and the sample was heated at 37°C for 1 hour, 70°C for 10 minutes and kept at 4°C until use. Cleavage reactions were electrophoresed using the Agilent 4200 TapeStation System (DNA 1000 kit, Agilent). The electrophoretogram profiles of fragment sizes resulting from sgRNA cleavage by Cas9 were analysed using the 2100 expert software package (Agilent). The editing efficiency was calculated as the area under the curve (AUC) for each cleaved fragment over the AUC of the uncleaved DNA fragment.

### RNP assembly and nucleofection

The 4D-Nucleofector™ X Unit (Lonza) was used for electroporation with the P3 Primary Cell 4D-Nucleofector™ X kit S for T cells and the SF Cell Line 4D-Nucleofector™ X kit S for HEK 293T cells. Firstly, 4 μg of TrueCutc Cas9 was precomplexed to 80 pmol of sgRNA (RNP) at room temperature for 20 minutes. 4 μM Alt-R^®^ Cas9 Electroporation Enhancer (IDT, 25 μM) was added to the RNP complex. T Cells were resuspended in P3 Primary Cell Nucleofector™ Solution and HEK293T cells were resuspended in SF buffer at 2×10^5^ cells/ 20 μl and the RNP complex was added immediately before transferring to a 100 μL Single Nucleocuvette for electroporation using program E0-115 for stimulated T cells or CM-130 for HEK293T cells. In HDR experiments 50 pmol of ssODN was added. Cells were recovered by adding pre-warmed (37°C) fresh media (without IL-2 supplementation) and incubated for 10 minutes at 37°C, 5% CO_2_. Cells were added to 500 μL of pre-warmed media with IL-2 (20 ng/mL) and IL-7 (2 ng/mL) in a 48 well dish and incubated for 72 hours (37°C, 5% CO_2_). In experiments including an Alt-R^®^ Cas9 HDR Enhancer (IDT), cells were added to 400 μL of media with IL-2 (20 ng/mL), IL-7 (2 ng/mL) and 30 μM of HDR enhancer in a 96 well plate and incubated for 12 hours (37°C, 5% CO_2_). 12 hours later, cells in media with the HDR enhancer were washed once with media and replaced with fresh media with IL-2 and IL-7 and incubated for a further 60 hours (37°C, 5% CO_2_).

### Cell viability

Cell viability was analysed before electroporation and before gDNA extraction with the LUNAC Automated Cell Counter using regular bright field counting (Logos Biosystems). The cell sample was mixed with acridine orange (Biotium, 10 ng/mL) at a ratio of 1:10. The viability was calculated as the percentage of live cells/mL to total cells/mL.

### gDNA extraction and Sanger sequencing

After 72 hours, gDNA was extracted using the DNeasy Blood & Tissue Kit (QIAGEN). The target region was PCR amplified using Phusion^®^ High-Fidelity PCR Kit (New England Biolabs) according to the manufacturer’s instructions with appropriate primers. The PCR products were electrophorised on a 1% agarose gel and PCR products were purified using QIAquick PCR Purification Kit (QIAGEN). Purified samples were sequenced by Sanger sequencing (Eurofins Genomics, SupremeRun Tube).

### Sequence deconvolution

SnapGene® was used to view and manipulate the sequencing files. Synthego ICE (Inference of CRISPR Edits) (www.ice.synthego.com) was used to analyse CRISPR edited samples. ICE compared the edited Sanger sequencing trace to an unedited control trace. Synthego ICE deconvolutes multiple edits created in a pool of sequences by separating out individual sequence traces and identifies the percentage of genomes that have indels. The software shows the sequences that are present in the edited population of cells and the percentage of sequences that have a particular edit.

### Immunoblotting

### Protein extraction

Cells were harvested 72 hours after electroporation, washed 3 times in PBS (1500 rcf for 5 minutes), resuspended in basal media free from IL-2 and IL-7 cytokines and incubated for 4 hours. Prior to lysis, cells were counted and stimulated in 100 μL of media with 20 ng/mL of recombinant mouse IL-6 (ELISA Std.) for 30 minutes (37°C, 5% CO_2_) in a 96 well plate. 2.5×10μ cells were used per sample. After stimulation cells were washed twice with PBS (1500 rcf for 5 minutes) and lysed with lysis buffer. The lysis buffer was made on the day and consisted of 1X Nupage™ LDS Sample Buffer (4×)(Invitrogen™) diluted with Milli-Q purified H_2_O, 10 μL/mL of Halt™ Protease and Phosphatase Inhibitor Cocktail (100×) (Thermo Scientific™), 0.2 M MgCl_2_ and 25 units/mL of lysate of Pierce™ universal Nuclease for Cell Lysis (Thermo Scientific™). Samples were centrifuged (2500rcf for 5 minutes) and the lysate was heated at 70°C for 10 minutes and kept at 4 °C.

### Gel Electrophoresis

SDS Running Buffer (20X) diluted with Milli-Q purified H_2_O at a ratio of 1:20 was prepared. The cell lysate was electrophoresed using Nu-Page 4-12% Bis-Tris gel. 18 μL of sample was loaded into each well. Precision Plus Protein™ Dual colour Standards molecular weight marker was used to identify protein sizes. The gel was run for 30 minutes at 60 v and 1 hour 50 minutes at 100 v.

### Membrane Transfer

Proteins were transferred from the gel onto a nitrocellulose membrane (Invitrogen™, iBlot™ 2 mini Transfer Stacks) using the iBlot 2 Dry Blotting System, program P0 (20 V for 1 minute, 23 V for 4 minutes, 25 V for 2 minutes).

### Membrane blocking

The membrane was blocked to prevent non-specific antibody binding in Odyssey^®^ Blocking Buffer (TBS) (LI-COR) and TBS (0.1% Tween) at a ratio of 1:2 and left for 1 hour on a rotor (fast).

### Antibodies

After blocking, the membrane was incubated for 1 hour with primary antibodies, STAT3 and pSTAT3. The membrane was washed twice with TBS (0.1% Tween) and incubated on the rocker for a further 30 minutes with secondary antibodies diluted in TBS (0.1% Tween) and Odyssey blocking buffer at a ratio of 1:1 (Table 3.2). The membrane was washed twice with TBS (0.1% Tween) and once with TBS prior to imaging.

Primary and secondary antibodies used for Western Blotting.

Primary and secondary antibodies used for Western Blotting
| Primary Antibody | Dilution | Secondary Antibody | Dilution |
| --- | --- | --- | --- |
| STAT3 (124H6) Mouse Anti-body<br>(Cell signalling) | 1:1000 | IRDye® 800CW Goat anti-Rabbit IgG (LI-COR) | 1:15,000 |
| Phospho-Stat3(Tyr705)(D3A7)<br>XP® Rabbit monoclonal Anti-body<br>(Cell signalling) | 1:1000 | IRDye® 680RD Goat anti-Mouse IgG (LI-COR) | 1:15,000 |

All antibodies were diluted in 10 mL of solution.

### LI-COR Odyssey CLx

The proteins were detected using infrared imaging on the Odyssey^®^ CLx Imaging System to detect 680 nm and 800 nm fluorescence on the same membrane. Image Studio™ Software was used for densitometry.

## Acknowledgments

Funding for the project was provided by the UK National Institute of Health Research Cambridge Biomedical Research Centre (NIHR BRC) (NIHR, grant number RG65966).

